# Changing dynamics of *Aedes aegypti* invasion and vector-borne disease risk for rural communities in the Peruvian Amazon

**DOI:** 10.1101/2024.09.04.611168

**Authors:** Kara Fikrig, Arnold O. Noriega, Rosa A. Rodriguez, John Bardales, José Rivas, Becker Reyna, Guido Izquierdo, Amy C. Morrison, Laura C. Harrington

## Abstract

*Aedes aegypti,* the primary vector of dengue virus, is predominantly considered an urban mosquito, especially in the Americas, where its reemergence began in cities after the end of continent-wide eradication campaigns. The results of our study diverge from this narrative, demonstrating the recent and widespread rural invasion of *Ae. aegypti* along major shipping routes in the northern Peruvian Amazon between the major cities of Iquitos, Pucallpa, and Yurimaguas. We identified *Ae. aegypti* populations in 29 of 30 sites surveyed across a rural to urban gradient and characterized mosquito larval habitats and *Ae. aegypti* adult metrics. Households, regardless of community size, were more likely to be positive for *Ae. aegypti* adult(s) and had a higher number of *Ae. aegypti* adults if a mosquito larval habitat was identified on the property, compared to houses without a larval habitat. In multiple instances, adult *Ae. aegypti* indices in rural villages were equal to or greater than indices in dengue-endemic cities, suggesting the entomological risk level in some rural areas is sufficient to sustain dengue transmission. Fourteen rural sites were sampled in transects from the community river port into town. In seven of these sites, houses closer to the port were significantly more likely to be infested with *Ae. aegypti* adults than houses further from the ports, and four additional sites had a marginal relationship to the same effect. This pattern suggests that many rural sites are invaded via adult *Ae. aegypti* disembarking from boats at the port, finding nearby oviposition sites, and advancing stepwise into town. The existence of the pattern also suggests that many of the sites are still experiencing active invasion, with sections of towns still *Ae. aegypti-*free. Only one site shows a strong signal of invasion via the egg or larval stage, with a focus of *Ae. aegypti* far removed from the port. The widespread infestation of *Ae. aegypti* in rural areas is a major public health threat given the far distance of communities to hospital care. It is important to implement control measures now before the mosquito gains a stronger foothold in zones of active invasion.

## Introduction

*Aedes aegypti* mosquitoes transmit numerous viruses impacting human health, including dengue virus, across the global tropics and subtropics. Dengue virus causes about 400 million infections per year, resulting in over 40,000 deaths [1, 2]. The disease can manifest a significant range of morbidity, including the need for advanced hospital care in the ICU, leading to a cumulative cost of about US$8.9 billion [1, 3].

Dengue was first reported in Peru in 1990, following closely on the heels of the reemergence of *Ae. aegypti* in the country. The mosquito was declared eradicated in Peru in 1958, as a result of the continent-wide Yellow Fever eradication campaign [4, 5]. In 1984, *Ae. aegypti* was detected once again in Iquitos, an Amazonian city only accessible by boat and plane [5]. In the span of three years, *Ae. aegypti* quickly reinvaded the city, increasing infestation levels from 1% to 26% by 1988 [5]. The first reported dengue outbreak in Iquitos occurred soon thereafter [5]. Similar patterns of *Ae. aegypti* eradication, reinfestation, increased dispersion, and dengue transmission repeated across much of the Americas during a similar time frame [6].

Because *Ae. aegypti* reinfestation and dengue transmission in the Americas began in densely populated urban areas, the vector and virus have been considered an urban problem [7]. Recently, the geographic profile of *Ae. aegypti* distribution and dengue transmission in the Americas has begun to change, expanding to more remote, rural communities [8]. There have been increasing reports of rural *Ae. aegypti* populations in numerous countries across the continent [9–13], which will likely expand further with increasing levels of transport connectivity, increasing use of plastics and other waste that can serve as larval habitat in rural areas, and persistent lack of access to running water, necessitating water collection [8]. This trend has received limited attention despite mirrored increases in rural dengue transmission, with evidence of high seroprevalence and the circulation of multiple serotypes in rural communities, particularly in Colombia and Ecuador [14–17]. Importantly, records of this rural expansion of the mosquito and virus is likely a substantial underestimate of their true rural distribution due to limited vector and disease surveillance in rural areas.

The northern Peruvian Amazon is an interesting setting to explore the expansion of *Ae. aegypti* given the heterogeneity of urbanization and fluvial connectivity throughout the region. Iquitos, the city where dengue was first identified in Peru, is connected by river to the port cities of Pucallpa and Yurimaguas, which also have endemic dengue transmission and established *Ae. aegypti* populations [18–20]. The rivers connecting these cities serve as a fluvial highway to transport goods and people between the cities, as well as the hundreds of rural communities scattered along the riverbanks.

The boats that travel these rivers play a critical role in *Ae. aegypti* mosquito dissemination to remote communities [21, 22]. A comprehensive survey of terrestrial and fluvial vehicles showed that all forms of fluvial transit could be infested with *Ae. aegypti,* with the highest infestation rate in large cargo boats (71.9%) [23]. Cargo boats were even found to sustain active oviposition by *Ae. aegypti,* allowing the mosquito to complete its full life cycle aboard the ship [24].

The first detection of *Ae. aegypti* in rural communities outside of Iquitos was in 2008, through an epidemiological study on arboviruses conducted by the U.S. Naval Medical Research Unit SOUTH (then known as NAMRU-6) and independent routine surveillance activities conducted by the Peruvian Ministry of Health [21]. In 2011-2012, a more comprehensive *Ae. aegypti* survey was conducted across 34 sites along the Amazon River and the 95 km stretch of road between Iquitos and Nauta (a small nearby city) [21]. These collections showed heterogeneous mosquito distributions in riverine communities and spatially explicit patterns along the roads. Half of surveyed riverine communities were infested. Communities with a larger human population size and closer distance to Iquitos were more likely to be infested with *Ae. aegypti.* Collections along the road showed presence of *Ae. aegypti* in all communities up to a discrete point, after which all further communities were negative. At the time, the furthest point of *Ae. aegypti* expansion from Iquitos was 37.1 km by river and 19.3 km by road [21].

In the intervening time, there has been no further systematic characterization of rural *Ae. aegypti* populations in the Peruvian Amazon, and no characterization of the mosquito beyond 95 km from Iquitos. The lack of information regarding vector movement and presence in remote areas is particularly concerning because communities in these areas are located far from medical facilities and care required to treat severe dengue. A combination of remote location and insufficient resources have created highly inequitable access to health care for rural communities in the Peruvian Amazon [25, 26].

Basic ecological information about the distribution of *Ae. aegypti* is vital to understand rural dengue risk in this region. To address this need, we conducted mosquito collections in thirty sites spread across a wide expanse of the northern Peruvian Amazon along over 1,000 km of river, across a rural – urban gradient, to determine the distribution of *Ae. aegypti* and measure the relative entomological risk for transmission of *Aedes-*borne viruses in the region. We expected to observe a trend in infestation levels associated with the degree of urbanization, anticipating that more populous communities would be more likely to be infested and exhibit higher entomological indices. Indeed, we did detect a trend of this nature, but it was weaker than expected because of the near ubiquity of *Ae. aegypti* across the transect. We detected *Ae. aegypti* infestations in almost all sites surveyed and measured high entomological indices in numerous small rural communities, suggesting that the expansion process is more advanced than previously imagined.

## Methods

In the following sections, we describe the rationale we used to select our sites, followed by the characteristics of the selected sites, mosquito collection methods, geographic data details, and the analysis methods. We then summarize our efforts to engage community members in the scientific process, and finish with statements on data availability, permits, and the availability of a Spanish translation of this paper.

### Site selection

In this study, we sought to characterize features of the invasion of *Ae. aegypti* into remote communities across a rural – urban gradient in the Loreto and Ucayali states in the Northern Peruvian Amazon. This region of the Amazon has several large and small cities and many towns and villages scattered throughout the immense expanse of rainforest. The communities are primarily connected by the complex matrix of rivers that cut through the forest. Mosquito collections were concentrated along the river system connecting Iquitos to Pucallpa, one of the two port cities that supply Iquitos with goods from outside the Amazon (Fig. 1), maximizing the likelihood of cargo ship-facilitated long-distance dispersal of *Ae. aegypti*.

**Figure 1.**
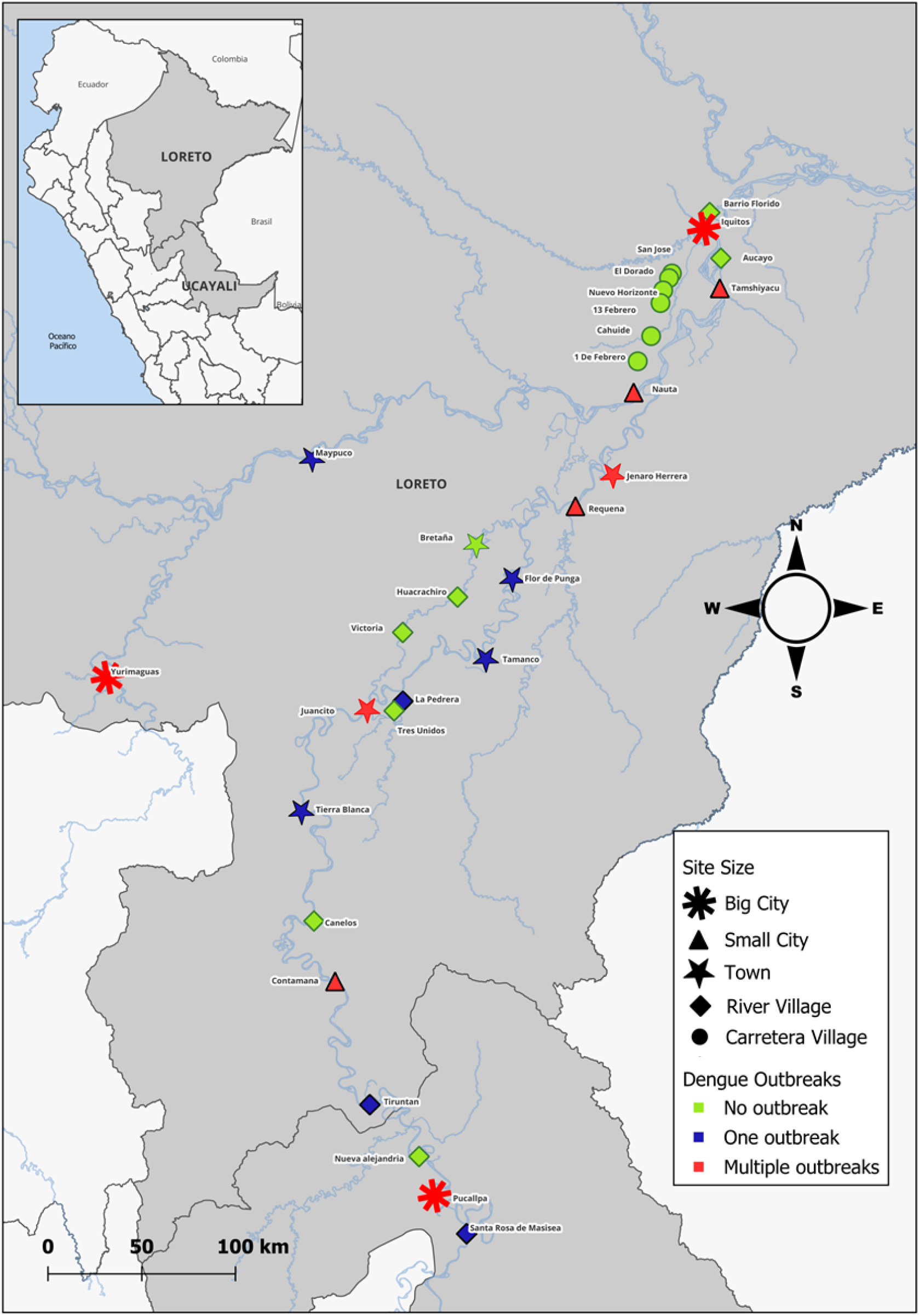
Map of sampling sites (30) across the regions of Loreto and Ucayali, demonstrating site size in symbols (black in legend) and history of dengue outbreaks in color (square in legend).

We aimed to select sites balanced across population size and incidence of known prior dengue outbreaks (none, one outbreak, or multiple outbreaks). Prior incidence of dengue was inferred from information shared by the Loreto health department about government-funded insecticide treatments in communities during the previous five years, record of which is synonymous with a reported dengue outbreak or malaria transmission. Malaria is not endemic in the selected communities, so insecticide treatment was in response to dengue outbreaks. Where there was no record of insecticide-use, there was no *a priori* record of *Ae. aegypti* presence.

Collections were conducted in two phases. The first phase was characterized by a higher number of household collections per site. Eighteen sites were selected between Iquitos and Pucallpa and one site on the far side of Pucallpa, roughly evenly spaced across the transect, as well as the two cities. In phase two, more limited collections were conducted on the short stretch of road that connects Iquitos to the small city of Nauta. Six sites were chosen, all of which were previously negative for *Ae. aegypti* in the 2011-12 collections by Guagliardo et al [21]. Collections were also conducted in one site beyond Iquitos down the Amazon River and one community on the Marañon river route to the other port city, Yurimaguas, as well as the city itself. In total, samples were collected in 30 sites, with 21 sites in phase 1 and 9 sites in phase 2.

### Site characteristics

The sites are geographically dispersed along two main fluvial transit routes: Iquitos – Pucallpa and Iquitos – Yurimaguas. The Iquitos – Pucallpa route spans a Euclidian distance of 536 km, which translates to 1,068 km and over 36 hours of travel on the fast boats that transit the winding Ucayali, Puinahua and Amazon rivers. The Iquitos – Yurimaguas route spans a Euclidian distance of 399 km, which translates to 661 km and over 20 hours of travel on the fast boats along the Huallaga and Marañon rivers. We also sampled two communities beyond the cities along these rivers. The population range of the sites is 120 to 484,000 people. Three sites are considered big cities (>23,000 people), four are small cities (5,000 – 23,000 people), seven are towns (1,000 to 4,999 people), and sixteen are villages (<1,000 people; ten along the rivers (= river village) and six along the Iquitos-Nauta highway (=road village)).

Among the twenty-one sites surveyed in phase 1 along the Iquitos-Pucallpa river route, seven communities never reported a dengue outbreak, six communities reported one dengue outbreak, and eight reported multiple outbreaks prior to the time of our collections (including Iquitos and Pucallpa, considered endemic for dengue). Five sites with one outbreak were initially selected as sites without a history of dengue (and no *a priori* knowledge of *Ae. aegypti* presence), but four experienced a dengue outbreak during the months between site selection and site visit and one experienced an outbreak just over five years prior, before the five-year time frame of health department data shared with our team. Among the nine sites surveyed in phase 2, seven communities had no record of locally transmitted dengue or *Ae. aegypti* (all the road villages and the river village beyond Iquitos), one had history of one outbreak, and one had multiple dengue outbreaks (Yurimaguas, also considered endemic).

While site urbanization level was classified by population size, other site characteristics are closely associated with these designations. The villages and towns had limited electrification, with electricity generated for about 3 - 4 hours per day (except for one town with full-day electricity), while the cities had full-day electricity. The houses in villages and towns tended to be made of wood boards, loosely fitted, with metal or palm leaf thatched roofs. In big cities, houses tended to be made of concrete with metal roofs, and small cities had a mixture of the two housing types. Most villages and towns did not have any form of official garbage management, except a few district capitals that had limited garbage collection and a dumping ground in the forest. Cities had more robust garbage management. While we consider villages and towns in this context to be rural, houses were notably aggregated, with agricultural lands dispersed along the river and into the forest. The yards were typically slightly larger than those in cities, but population density is not as dramatically different as rural/urban dichotomies in other regions.

### Mosquito Collection

Mosquito collections were conducted with Prokopack aspirators [27] operated inside structures. Most collections were conducted in residences, and infrequently inside stores, offices, and structures of mixed used, with a store or office space alongside a living space. Structures were searched exhaustively in all spaces where collectors were permitted to enter. Mosquitoes were maintained alive in mesh-covered 0.7L collection cups, labeled with the unique structure code. Mosquitoes were killed at the end of the day by placing cups inside a sealed bag with acetone for approximately 15 min. Specimens were sorted and all *Ae. aegypti* were confirmed based on key taxonomic features with a field microscope [28].

In addition, larval surveys were conducted in and outside of a subset of structures where adult collections were performed, except for Iquitos, where no larval inspections were conducted. It was noted whether a property had at least one container with live mosquito larvae or pupae, regardless of mosquito species. Time constraints prevented conducting more comprehensive larval surveys across all properties and containers.

In phase 1 collections (along the Iquitos-Pucallpa transect), the sampling approach was designed to include areas near and far from the ports and was adapted based on community size and boat schedules. In villages, a high percentage of houses were visited (37 – 93% of houses based on 2017 census household counts [29, 30]) and the majority approached granted entry to our team. In towns, we collected in areas near and far from community ports; whenever possible, we collected in a transect from the town port to residences farthest away from the river. In towns, we sampled 12 – 31 % of houses. In cities, we also collected close to and far from the port(s). Notably, city collections were the least representative of the entire city. The proportion of houses surveyed was low (< 0.2% of houses) and the sampling area was influenced by health department recommendations regarding where there was high *Ae. aegypti* prevalence.

In phase two (communities along the Iquitos-Nauta road, the Yurimaguas river route, and downriver from Iquitos), collections were limited to a smaller number of houses (13 – 40 houses; coverage ranging between 13 – 70% for the villages, 8 – 13% for the towns, and 0.1% for the city, Yurimaguas).

We sampled from January through June 2023, with most collections corresponding to the period of higher rainfall, river levels, and dengue transmission.

Depending on structure size and other logistical considerations, collections were sometimes conducted by one collector, sometimes by two collectors, and rarely by three or four collectors. Five different collectors were involved throughout the study, but only three or four collectors were involved in the collections for any given site.

### Geographic Data Collection

At the time of sampling, a GPS point was taken at the front door of the property using the UTM Geo Map application (Y2 Tech, Indonesia). The GPS point was taken when the application-reported accuracy fell within 3m. The GPS coordinates were uploaded to qGIS (version 3.30) and overlaid with 2023 Bing Maps satellite images. The Euclidian distance was measured between each sampling point and the community port (identified via images taken during rainy season). In some communities, there are two separate ports for rainy and dry season, or for barges and passenger boats. In these cases, we selected the distance to the closest of the two ports for each sampling point for subsequent analyses.

### Data Analysis

All analyses were performed in R version 4.3.0 [31].

#### Adult *Aedes aegypti* Infestation levels

For each community, the following metrics were calculated: adult house index (percentage of houses infested with adult *Ae. aegypti;* **AHI**) and mean adult number (average number of *Ae. aegypti* adults per house). Maps displaying infestation levels across the region were created using qGIS.

#### Impact of Urbanization level on Adult *Ae. aegypti* Infestation Levels

Generalized linear mixed models (GLMM) were utilized to determine the impact of urbanization level (i.e. site size) on infestation metrics (presence of adult *Ae. aegypti* and number of adult *Ae. aegypti* per house). Post hoc pairwise comparisons were conducted with the ‘emmeans’ and ‘pairs’ functions with Bonferroni adjustment using the ‘emmeans’ package [32] to compare the estimated marginal means of the infestation metrics for each urbanization level, adjusting for other variables in the model. Urbanization was considered an ordinal categorical variable with five levels of increasing population size: road village, river village, town, small city, and big city. Notably, the road villages are not uniformly smaller than river villages but had a lower population on average and were grouped separately due to the distinct ecology of their locations.

##### Presence of adult Ae. aegypti

A GLMM was conducted to determine the impact of urbanization level on the probability of whether a house was negative (0) or positive (1) for at least one *Ae. aegypti* adult using the lme4 package [33] with a binomial distribution. The fixed effect was the urbanization level, and the random effects included the site, the site nested within urbanization level, and the collector team (each unique collection team, whether an individual or combination of individuals, was considered a separate level of the variable).

##### Number of adult Ae. aegypti

Another GLMM was performed to determine the relationship between the urbanization level and number of adult *Ae. aegypti* per house. Due to overdispersion of the data, a glmmTMB model with a negative binomial distribution was used [34], with the same fixed and random effects as above.

Larval Infestation Levels and Impact of Larval Habitat on Adult Infestation Levels.

The larval house index was calculated (percentage of houses with at least one mosquito larval habitat with live larvae or pupae). Generalized linear mixed models were then used to examine the relationship between the presence of a mosquito larval habitat on a property and the adult *Ae. aegypti* infestation levels. These models were conducted for the subset of data for which larval habitat data was collected. To further explore the interaction between larval habitat and urbanization level, the ‘emtrends’ function from the ‘emmeans’ package [32] was used to estimate the effect of larval habitat across different urbanization levels.

##### Presence of adult Ae. aegypti

A GLMM with a binomial distribution was conducted to determine the relationship between the presence of a mosquito larval habitat and the presence of at least one adult *Ae. aegypti* mosquito for each house. The fixed effects included presence of a larval habitat and urbanization level, and the interaction between the two. The random effects included the site nested within the urbanization level and the collector team.

##### Number of adult *Ae. aegypti*

Another GLMM was performed to determine the relationship between the presence of a mosquito larval habitat and number of adult *Ae. aegypti* per house using a glmmTMB model with a negative binomial distribution, with the same fixed and random effects as above.

#### Impact of distance to port on infestation levels

Generalized linear mixed models were performed on a subset of communities to better understand the impact of distance to the community port on *Ae. aegypti* infestation. Analyses were limited to riverine communities with a minimum of 60 data points, where at least 10% of houses were sampled (based on number of households in the 2017 census [29, 30]), and where we conducted transects leading away from the port. This subset included 14 communities. To assess how the effect of distance from port varies by community, we used the ‘emtrends’ function [32] to estimate this effect for each community.

##### Presence of adult Ae. aegypti

First, we performed a GLMM with a binomial distribution to evaluate the impact of distance from the port on whether a house was positive for at least one *Ae. aegypti* adult. Fixed effects included the site, distance (km) of the house to the port, and the interaction of the two. The collector team was included as a random effect. The BOBYQA optimizer with 200,000 iterations was utilized.

##### Number of adult Ae. aegypti

We then attempted to follow a similar methodology to determine the impact of distance to the port on the number of adult mosquitoes per house, using glmmTMB with a negative binomial distribution [34]. The model failed to converge. Subsequently, the data was restricted to the non-zero values, to determine if there was an impact of distance to port on the number of *Ae. aegypti* in a house, among those houses that were infested with at least one *Ae. aegypti* adult. A glmmTMB model with a truncated Poisson (link=log) distribution was used with the same fixed and random effects as in the presence model.

##### Community Engagement

Our team took several steps to engage communities in the research. Before beginning collections, we asked permission from the leadership of each location. In cities and larger towns, we met with the environmental health leadership in the respective health department. In villages and smaller towns, we asked permission from elected government officials and, in communities with traditional indigenous leadership, from the community Apu. The collections were conducted on the ancestral and current lands of several indigenous peoples, including the Cocama-Cocamilla, Shipibo-Conibo, and Urarinas peoples. Once we received permissions from the corresponding leadership, we proceeded to ask individual permission from each household. After each household visit, we discussed our findings with household members and explained how to prevent future larval habitats, if we found any.

In many of the towns and villages, we conducted a workshop about *Ae. aegypti* and dengue and shared details about the scientific purpose of our visit. The workshops were hosted for different audiences depending on the circumstances, including the general community, primary or secondary school children, or the health department employees.

We recruited one representative from each of twenty sites with whom we have maintained contact, shared results, and provided opportunities to ask questions and give feedback. Most of the communities were also visited at the conclusion of the study after the results were analyzed to share results in meetings with the full community, the municipality, and/or the health center employees, depending on the circumstances. All results were also shared in detail with the regional health departments.

### Ethics Statement

All necessary permits and approvals were obtained, including the SERFOR collection permit (RD N°000066-2024-MIDAGRI-SERFOR-DGGSPFFS-DGSPFS), Gerencia Regional de Salud approval (N° 256-2022-GRL-DRSL/30.09-INVESTIGACIÓN), and household collections were deemed exempt by the Naval Medical Research Unit SOUTH IRB (NAMRU6.P0003) and the Cornell University IRB (IRB0145012). Formal verbal consent was obtained from the communities’ leadership as well as from each engaged household, which is detailed in the *Community Engagement* subsection.

### Data Availability

All data and code will be deposited in Cornell University Library’s institutional repository, eCommons (https://ecommons.cornell.edu) for preservation and access without restriction.

### Spanish Translation

This paper has been translated into Spanish to improve accessibility in the region where the research was conducted (see Supplemental File 1).

## Results

### Adult Ae. aegypti infestation levels

We observed a wide range of infestation levels (Fig. 2; Table 1). Among the large cities, Iquitos had both the highest adult house index (percent of houses with at least one adult *Ae. aegypti;* AHI = 91.7%) and highest mean number of adult *Ae. aegypti* per house (mean ± SD = 7.67 ± 9.46). Yurimaguas had an AHI of 83.3% and mean adult number of 2.29 ± 2.05, while Pucallpa had an AHI of 52.9% and mean adult number of 1.83 ± 3.71.

**Figure 2.**
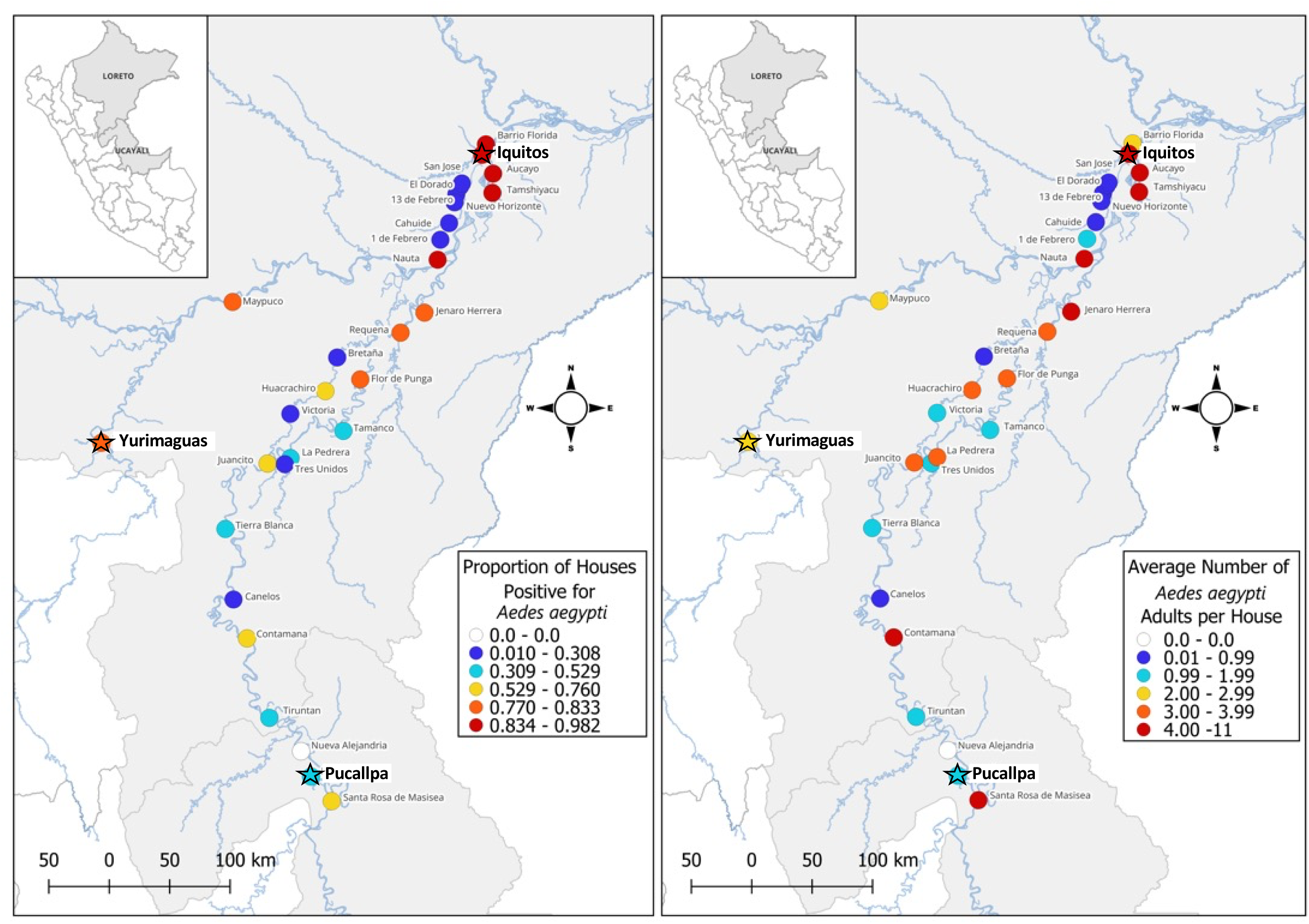
Maps of adult *Ae. aegypti* infestation levels in our sampling sites across the regions of Loreto and Ucayali in the Peruvian Amazon. On the left, the point color represents the proportion of houses positive for *Ae. aegypti,* with white representing absence, and from blue to red representing an increasingly higher proportion of houses positive for the mosquito. On the right, the same sites are colored by the average number of adult *Ae. aegypti* collected per house, again with white representing zero *Ae. aegypti* and from blue to red representing an increasingly higher average number of adult *Ae. aegypti* per house. The stars highlight the three major cities.

**Table 1.** Infestation levels of adult *Ae. aegypti* and larval mosquitoes by site and urbanization level.

| Site | Pop. Size<sup>§</sup> | No. Houses Visited | Adult *Ae. aegypti* House Index (%) | Mean No. Adult *Ae. aegypti* (± SD) | No. Houses Larval Survey | Larval Index (%) |
| --- | --- | --- | --- | --- | --- | --- |
| Large City |  |  |  |  |  |  |
| Iquitos | 437,620 | 48 | 91.7 | 7.67 (9.46) | 0 | NA |
| Pucallpa | 310,750 | 140 | 52.9 | 1.83 (3.71) | 138 | 16.7 |
| Yurimaguas | 41,827 | 24 | 83.3 | 2.29 (2.05) | 24 | 33.3 |
| Small City |  |  |  |  |  |  |
| Contamana | 17,429 | 76 | 75.0 | 5.21 (8.93) | 68 | 32.4 |
| Nauta | 19,551 | 67 | 88.1 | 5.16 (5.10) | 28 | 46.4 |
| Requena | 22,875 | 97 | 79.4 | 3.76 (6.14) | 28 | 28.6 |
| Tamshiyacu | 6,181 | 57 | 98.2 | 11.00 (12.84) | 30 | 63.3 |
| Town |  |  |  |  |  |  |
| Bretaña * | 1,686 | 132 | 21.2 | 0.74 (2.66) | 109 | 14.7 |
| Flor de Punga | 1,763 | 78 | 79.2 | 3.77 (3.74) | 72 | 43.1 |
| Jenaro Herrera | 3,596 | 77 | 83.1 | 4.97 (8.20) | 26 | 42.3 |
| Juancito | 2,124 | 100 | 76.0 | 3.04 (4.46) | 62 | 41.9 |
| Maypuco | 1,248 | 24 | 83.3 | 2.58 (1.98) | 24 | 37.5 |
| Tamanco † | 1,738 | 162 | 40.7 | 1.56 (3.26) | 154 | 29.9 |
| Tierra Blanca † | 1,602 | 112 | 41.1 | 1.63 (3.51) | 109 | 36.7 |
| Riverine Village |  |  |  |  |  |  |
| Aucayo | 587 | 65 | 84.6 | 8.48 (9.79) | 29 | 58.6 |
| Barrio Florida | 673 | 26 | 84.6 | 2.38 (2.80) | 26 | 46.2 |
| Canelos * | 317 | 60 | 23.3 | 0.72 (1.98) | 59 | 32.2 |
| Huacrachiro * | 667 | 87 | 62.1 | 3.15 (4.52) | 55 | 41.8 |
| La Pedrera | 531 | 65 | 38.5 | 3.62 (9.83) | 63 | 31.7 |
| Nuevo Paris / Nueva Alejandría | 361 | 45 | 0 | 0 | 45 | 0 |
| Santa Rosa de Masisea † | 413 | 85 | 68.2 | 4.05 (7.48) | 83 | 34.9 |
| Tiruntan † | 640 | 99 | 40.4 | 1.30 (3.12) | 99 | 32.3 |
| Tres Unidos | 507 | 62 | 24.2 | 1.39 (4.36) | 61 | 27.9 |
| Victoria * | 858 | 118 | 19.5 | 1.00 (3.30) | 29 | 20.7 |
| Road Village |  |  |  |  |  |  |
| 1 de Febrero * | 183 | 13 | 30.8 | 1.23 (2.80) | 13 | 15.4 |
| 13 de Febrero * | 670 | 36 | 22.2 | 0.89 (2.05) | 24 | 12.5 |
| Cahuide * | 794 | 40 | 17.5 | 0.35 (0.92) | 40 | 5.0 |
| El Dorado * | 90 | 21 | 14.3 | 0.19 (0.51) | 21 | 4.8 |
| Nuevo Horizonte * | 308 | 21 | 9.5 | 0.14 (0.48) | 21 | 9.5 |
| San Jose * | 123 | 20 | 10.0 | 0.15 (0.49) | 20 | 10.0 |

‡ Big city data [35] and other site data [36, 37]
\* first report of *Ae. aegypti* presence in the community
† first dengue outbreak occurred in the community during the year of collections

Among the four small cities, Tamshiyacu had the highest adult house index and mean adult number (AHI= 98.2%; mean = 11.0 ± 12.8), even surpassing the infestation levels in Iquitos. The other three small cities had an AHI between 75.0% and 88.1% and mean number of adults between 3.76 and 5.21.

Among the seven towns, the AHI ranged from 21.2% in Bretaña to 83.3% in Maypuco. The town with the lowest mean number of adults was also Bretaña (0.74 ± 2.66) and the town with the highest mean number of adults was Jenaro Herrera (4.97 ± 8.20).

Among all 30 sites, only one did not have *Ae. aegypti* – Nueva Alejandría/Nuevo Paris, two small riverine villages next to one another and connected by road, which we considered as one site. The other nine riverine villages all had *Ae. aegypti*, with different levels of infestations. Some were highly infested, such as Aucayo (AHI= 84.6%; mean=8.48 ± 9.78) and Santa Rosa de Masisea (AHI= 68.2%; mean=4.05 ± 7.47), while others had low levels of infestation, such as Victoria (AHI= 19.5%; mean=1 ± 3.30) and Canelos (AHI= 23.3%; mean = 0.72 ± 1.98).

All six road villages had *Ae. aegypti* infestations, but at relatively low levels, with the AHI ranging from 10.0% in Nuevo Horizonte to 30.8% in 1 de Febrero and the mean number of adult *Ae. aegypti* ranging from 0.14 ± 0.48 to 1.23 ± 2.80, respectively (Fig. 3).

**Figure 3.**
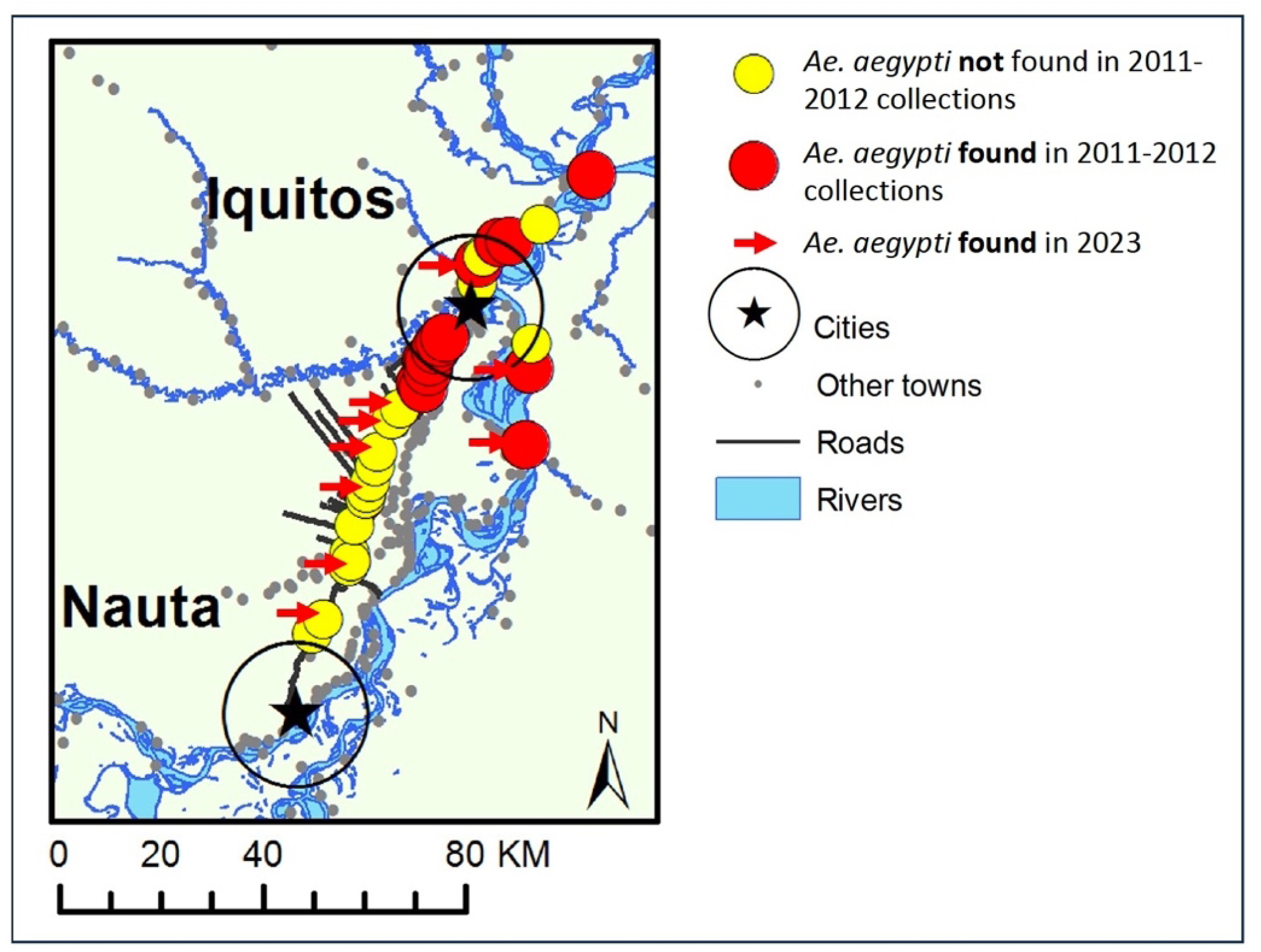
Adapted from Guagliardo et al. (2014) [21]. The map shows the original sampling sites from 2011-2012 conducted within 95 km of Iquitos. Red circles show sites where *Ae. aegypti* was found to be present and yellow circles show where it was absent. Red arrows show the sites where collections were conducted for the current study in 2023, all of which were positive for *Ae. aegypti*.

*Aedes aegypti* was reported for the first time at ten sites, including four on the river (Bretaña, Canelos, Huacrachiro, and Victoria) and six on the Iquitos-Nauta road (1 de Febrero, 13 de Febrero, Cahuide, El Dorado, Nuevo Horizonte, and San Jose). An additional four communities on the river would have been first reports at the time of site selection, but dengue outbreaks occurred in the months immediately prior to our visit, alerting authorities to the presence of the vector (Tamanco, Tiruntan, Tierra Blanca, and Santa Rosa de Masisea).

### Impact of Urbanization Level on Adult Ae. aegypti Infestation Levels

#### Presence of adult *Ae. aegypti*

Urbanization level had a positive linear relationship (p<0.0001) and a marginally significant negative quadratic relationship with *Ae. aegypti* adult presence in a house (p=0.072). In other words, the presence of *Ae. aegypti* adults in houses initially increases with urbanization level, but the quadratic term suggests that the probability of *Ae. aegypti* presence may decline at higher urbanization levels, indicating a more complex, nonlinear pattern. This relationship can be better understood through pairwise comparisons of the estimated marginal means at each urbanization level. The only significant differences between urbanization levels were lower infestation levels in road villages compared to towns (p=0.006), small (p<0.001) and big cities (p=0.003), as well as lower infestation levels in river villages compared to small cities (p=0.008). The trend can also be visualized in the raw data, which indicates an increase and then plateau of AHI with increasing urbanization (Fig. 4A).

**Figure 4.**
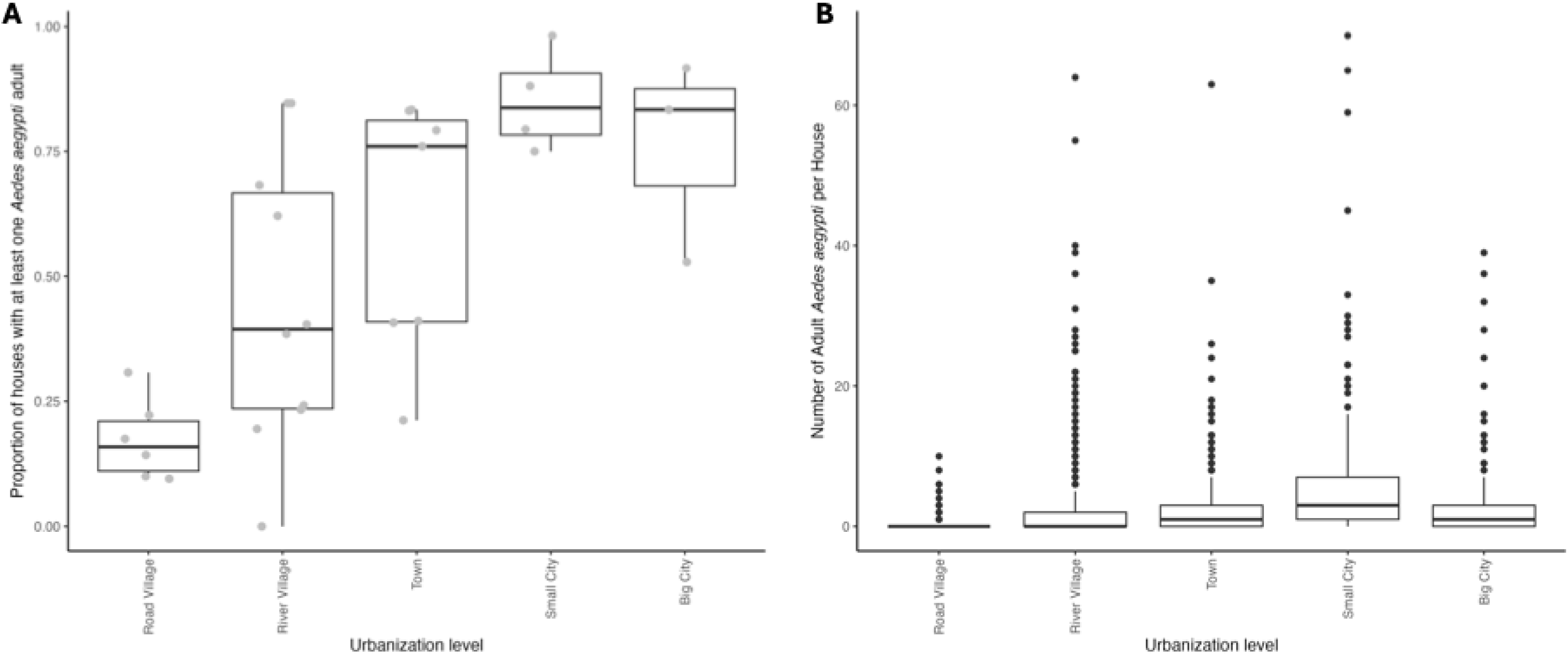
A) The box plot shows the proportion of houses with at least one *Ae. aegypti* adult per site, aggregated by urbanization level. The overlaid gray points display raw data for each site. **B)** The box plot displays the number of *Ae. aegypti* adults collected per house, aggregated by urbanization level. Notably, there are many outliers, demonstrating that houses at every urbanization level can yield extremely high numbers of adult *Ae. aegypti,* except for road villages, which had fewer and less dramatic outliers.

#### Number of adult *Ae. aegypti*

In the case of the number of *Ae. aegypti* adults per house, the model revealed a positive linear relationship (p<0.0001) and a negative quadratic relationship with urbanization level (p=0.028). The pairwise comparisons closely mirrored the household positivity results, with road villages having significantly lower numbers of *Ae. aegypti* per house compared to towns, small, and big cities (p<0.010) and river villages with significantly lower numbers compared to small cities (p=0.027). The trend can also be visualized in the raw data, with an increase in the number of mosquitoes per house with urbanization level until a drop in numbers from small to big cities (Fig. 4B).

### Larval Infestation Levels and Impact of Larval Habitat on Adult Infestation Levels

Most communities had a lower percentage of houses positive for mosquito larvae than the percentage positive for adult *Ae. aegypti*, with the exceptions of Canelos, Tres Unidos, and Victoria, which had a higher percentage positive for mosquito larvae, and Nuevo Horizonte and San Jose, which had the same percentage positive for mosquito larvae and adult *Ae. aegypti.* Notably, all these sites have a low burden of *Ae. aegypti* infestation in general (larval index between 9.5% – 32.2% and AHI between 9.5% – 24.2%).

#### Presence of adult *Ae. aegypti*

Properties with a larval site were more likely to have at least one adult *Ae. aegypti* in the house, regardless of site size (GLMM, p < 0.01). In big cities, 83.9% of houses with a larval habitat were positive for adult *Ae. aegypti*, whereas 51.1% of houses without a larval habitat were positive for adult *Ae. aegypti.* In small cities, these numbers were 93.4% in houses with larval habitat and 73.9% in houses without larval habitat; towns 71.5% and 39.9%; river villages 74.3% and 32.2%; and road villages 83.3% and 11.0%, respectively.

#### Number of adult *Ae. aegypti*

Similarly, the mean count of adult *Ae. aegypti* per house was higher in houses with a larval habitat compared to houses without a larval habitat (GLMM, p < 0.001). In big cities, houses with a larval habitat had an average of 4.5 ± 6.1 adult *Ae. aegypti* per house, whereas houses without a larval habitat had an average of 1.2 ± 1.8 adults. In small cities, houses with a larval habitat had 8.1 ± 12.2 adults while houses without a larval habitat had 3.5 ± 4.4; in towns 3.8 ± 4.5 and 1.3 ± 3.3, respectively; in river villages, 5.3 ± 8.2 and 1.0 ± 2.8, respectively; and in road villages, 2.6 ± 2.8 and 0.3 ± 0.9, respectively.

### Distance from port

In the riverine towns and villages where we sampled in transects from the port, we repeatedly found *Ae. aegypti* in the houses closer to the ports more often than in houses further from the port (Fig. 5 and 6).

**Figure 5.**
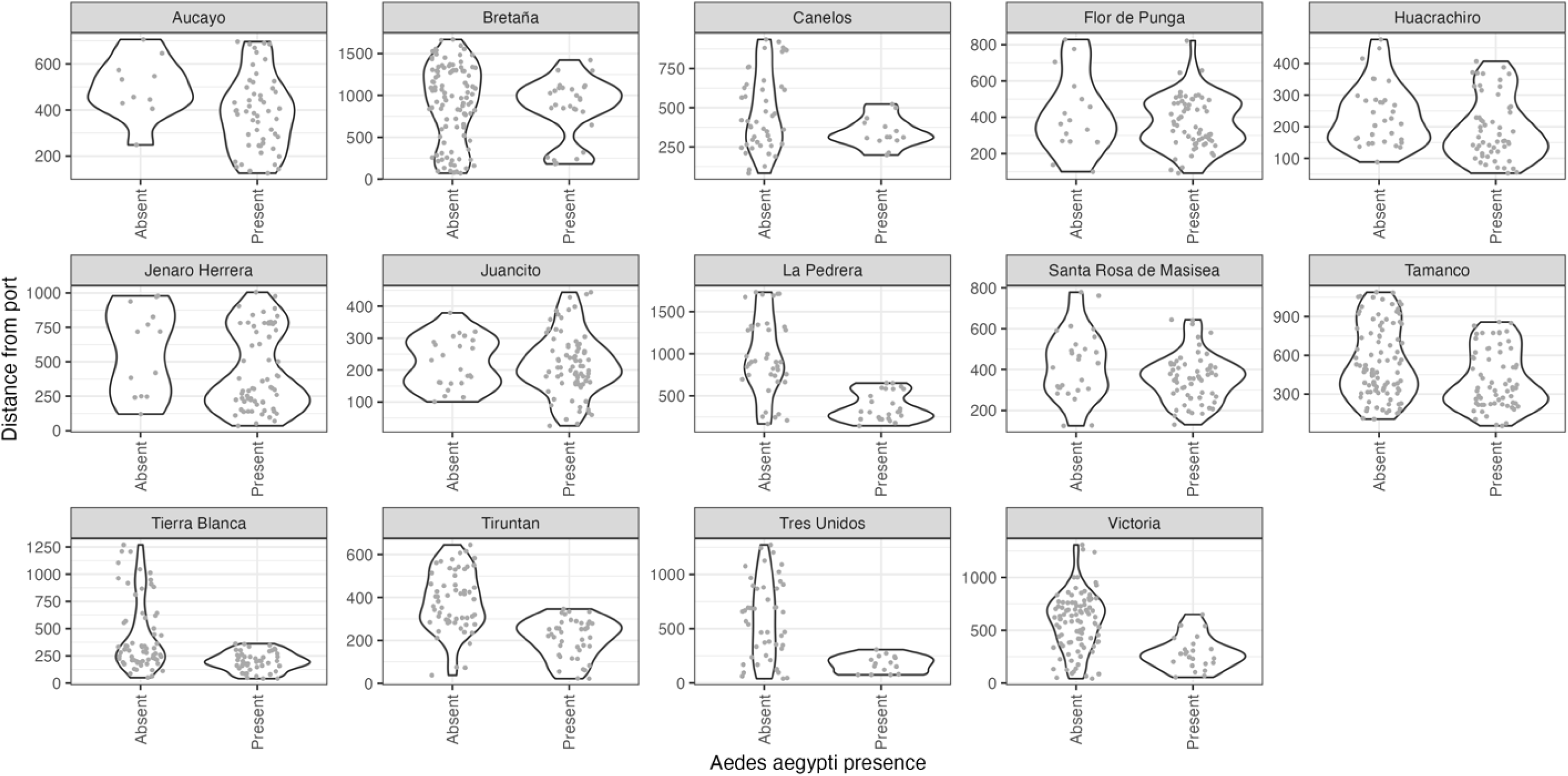
The violin plots display the distance of each household mosquito collection from the community port and whether *Ae. aegypti* was absent or present at the house. Each facet displays the results for a separate riverine village or town, limited to those sites where collections were conducted in transects from the port. The shape of the violin plot demonstrates the density of *Ae. aegypti-*positive and -negative households across the distances sampled. A repeated pattern can be observed: houses without *Ae. aegypti* tended to be concentrated further from the port.

**Figure 6.**
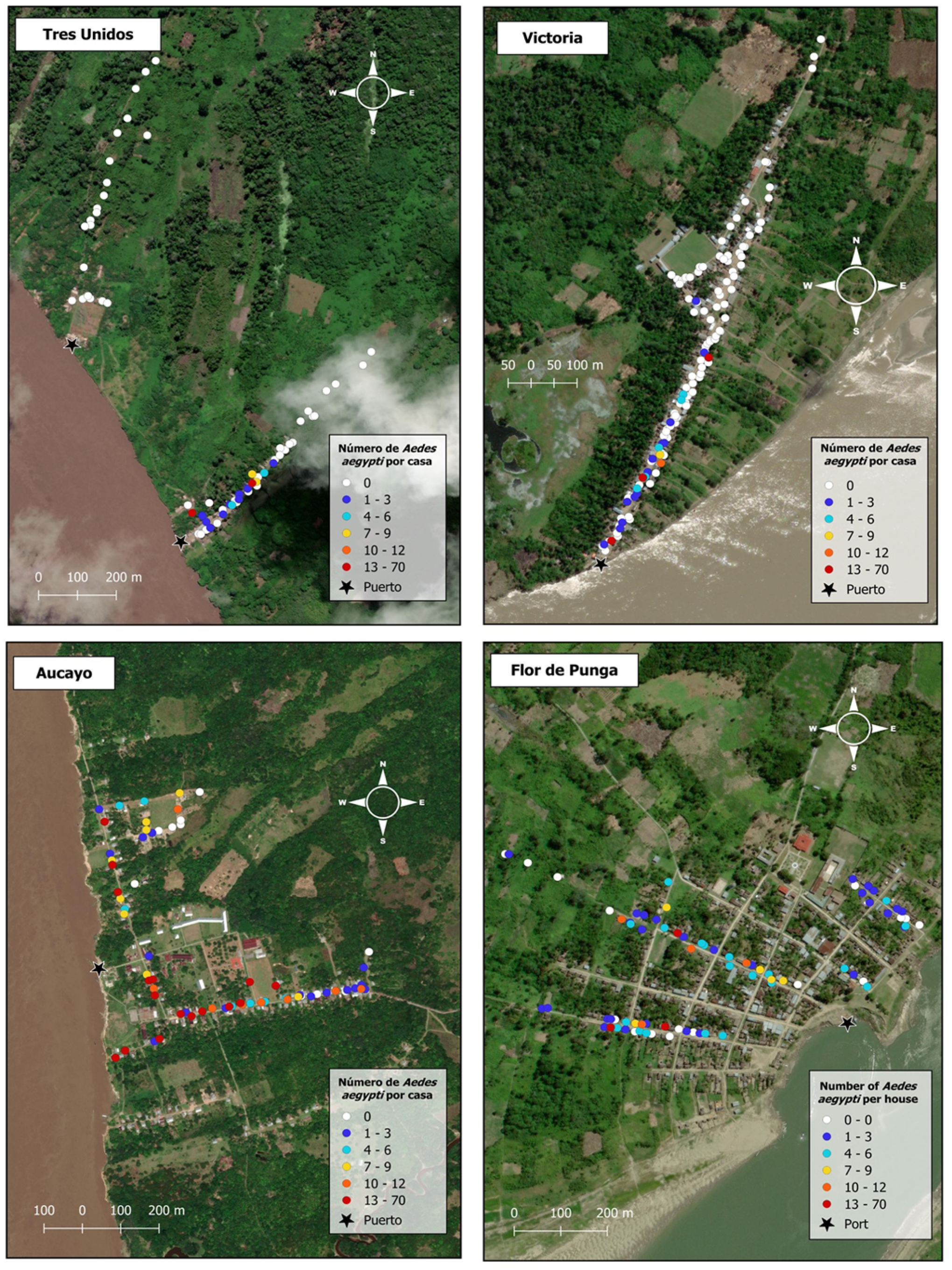
Maps of four sites (3 river villages and 1 town – Flor de Punga). Each point represents a household collection event. The color indicates the number of *Ae. aegypti* collected in the house. White points represent houses where the mosquito was absent and colors from blue to red indicate an increasing number of *Ae. aegypti* per house. In Tres Unidos, the village is divided into two neighborhoods due to landslides and erosion resulting sections of riverbank falling into the river (“desbarrancamiento”), forcing people to move over the past 5 years. The new neighborhood was *Ae. aegypti*-free, while the old neighborhood had *Ae. aegypti* concentrated near the port. Similarly, in Victoria, *Ae. aegypti* was present near the port and disappeared at a certain point into the town. In contrast, *Ae. aegypti* was highly dispersed in Aucayo, although most of the negative houses were located farthest from the port. In Flor de Punga, *Ae. aegypti* was also dispersed throughout the town, with the negative houses scattered without a clear pattern. Additional community-level maps can be found in Supplemental Fig. 1.

#### Aedes aegypti presence

The observed mosquito distribution pattern was interrogated with a GLMM to determine the relationship between distance to port and probability of *Ae. aegypti* presence (Fig. 7). In seven communities, houses closer to the port were more likely to be *Ae. aegypti*-positive than houses further from the port (p < 0.05). Four communities had a marginal relationship between distance and probability of *Ae. aegypti* presence (0.05 ≤ p < 0.10). Three communities had no relationship between distance to port and the likelihood of *Ae. aegypti* presence.

**Figure 7.**
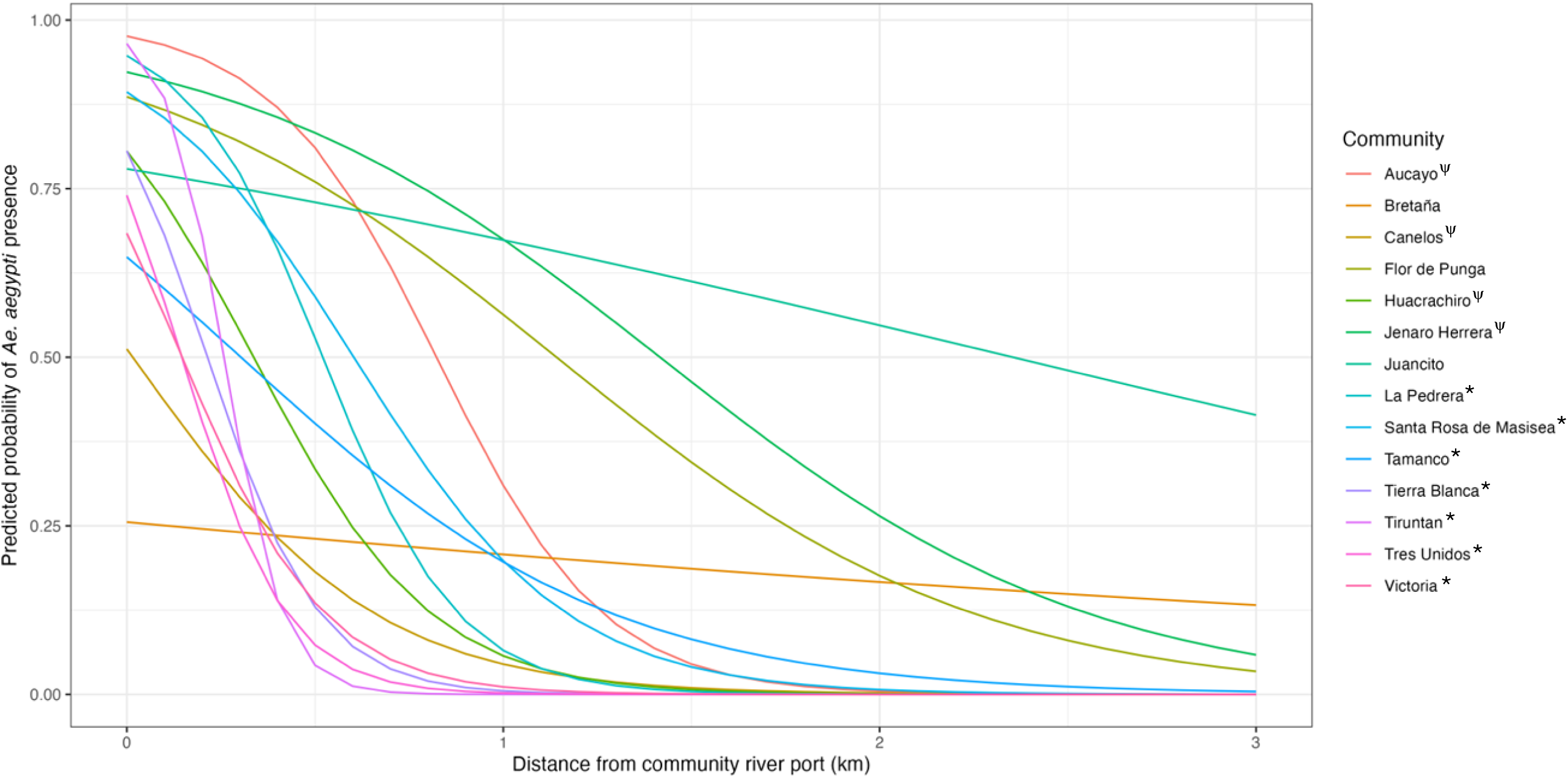
The output of the GLMM supports a repeated relationship between the predicted probability of *Ae. aegypti* presence in a house and the distance of the house from the community port. Each line represents the relationship for each of the 14 communities included in the analysis. In the graph legend, the * represents communities with a significant relationship between presence and distance (7 communities with p < 0.05) and ψ represents communities with a marginal relationship (4 communities with 0.05 ≤ p < 0.1).

#### Aedes aegypti abundance

A glmmTMB model with a negative binomial distribution was used to determine whether a similar relationship existed between distance from the port and the number of adult mosquitoes in the house. However, the model failed to converge due to a non-positive-definite Hessian Matrix. Data was restricted to non-zero values and the analysis repeated to estimate the effect of distance from the port on the number of *Ae. aegypti* in houses that had at least one *Ae. aegypti* mosquito. No clear trend emerged – eight of fourteen communities had no significant relationship between distance and number of *Ae. aegypti,* four communities had a significant decrease in the number of mosquitoes further from the port, and two had a significant increase in mosquitoes in houses further to the port.

## Discussion

Our study demonstrates that *Ae. aegypti* has invaded rural communities throughout a large region of the Peruvian Amazon, including numerous remote villages with no previous report of this vector. Our results also suggest a mechanism of invasion – adult *Ae. aegypti* flying off boats docked at community ports. The extent and ubiquity of *Ae. aegypti* invasion is alarming and poses a growing public health threat for numerous vector-borne diseases in rural Amazonia.

Our collections show that distance from a major city does not limit *Ae. aegypti* infestation along major shipping routes. The vector was found at the furthest site sampled, Victoria, 266 km from the closest city, Iquitos. The only site where *Ae. aegypti* was absent was a pair of communities near Pucallpa. This widespread dispersal, independent of distance from a major city, differs from the pattern observed in 2011-2012, when the first systematic study of rural *Ae. aegypti* expansion in Peru was conducted in communities within a 95 km radius of Iquitos [21]. At the time, communities with *Ae. aegypti* tended to be closer to Iquitos than *Ae. aegypti-*negative communities. This change may reflect the vector’s expansion to more remote areas since the original study, neutralizing this pattern. Alternatively, the observed pattern may have been an artifact of the specific sites selected, and invasion to sites further from Iquitos may have already occurred at the time of the 2011-2012 study.

Our collections also show a dramatic expansion of *Ae. aegypti* in road village sites along the Iquitos-Nauta highway. All six villages sampled were free of *Ae. aegypti* during collections conducted in 2011-2012. By 2023, all six communities were infested with *Ae. aegypti,* spanning the full length of the 95 km highway.

The level of *Ae. aegypti* infestation in the rural villages, towns, and small cities was surprising. We documented ten first reports of the mosquito in rural communities and towns. While there was a trend of increasing infestation levels with increasing urbanization, this relationship was not perfectly linear and appears to be driven by the particularly low infestation levels in road villages, as well as the particularly high infestation levels in small cities. There was a remarkable level of variation in infestation levels between sites within the same site size category. It is also notable that *Ae. aegypti* larval and adult indices observed in our study far exceeded those reported in previous studies carried out in Iquitos, Peru, with mean *Ae. aegypti* collected per house rarely exceeding 1 and larval house indices less than 45% [38–41]. At some rural sites, the percentage of houses positive for adult *Ae. aegypti* and the average number of *Ae. aegypti* per house surpassed infestation levels in the large cities, demonstrating that entomological risk levels in some rural areas are sufficient to sustain a dengue outbreak. This is particularly striking because sampling locations in big cities was influenced by health department recommendations regarding where there was high *Ae. aegypti* prevalence, which may have biased the results in cities towards higher infestation levels than would be found through random sampling.

Within rural villages and towns, we repeatedly observed that *Aedes aegypti* adults were significantly more likely to be found in houses closer to the community port compared to houses further from port in seven of fourteen communities, with a marginally significant relationship in four additional communities. This result supports previous research in the region attributing long-distance dispersal to *Ae. aegypti* infestation on boats [23, 24]. It also adds a layer of detail – the invasions most likely occur at the adult life stage, not the egg or immature life stages. The higher probability of presence near ports suggests that adult female *Ae. aegypti* fly off boats, oviposit in nearby houses, and subsequent generations advance stepwise into the community, creating a distinct distribution pattern radiating from the port. If invasion occurred in the egg or immature stages, we would expect a more heterogenous mosquito distribution, as infested containers should be equally likely to be brought to any house in the community from the boat. Although we believe the distribution patterns indicate adult-stage invasion, it is not conclusive proof and may have another explanation.

There were a few exceptions to this trend. In Juancito and Flor de Punga, which had no relationship between distance and *Ae. aegypti* presence, the species was widespread throughout the town, with an AHI over 75%. The mosquito likely established earlier at these sites and completed its invasion into the towns before we sampled. Three of the four communities with a marginally significant relationship between *Ae. aegypti* presence and distance to the port also had relatively widespread adult presence (greater than 60% AHI), suggesting that the weakening impact of distance to port may also be due to *Ae. aegypti* reaching establishment. This is further supported by prior evidence that Aucayo village was already infested with *Ae. aegypti* during 2008 and 2012 collections [21]. The only low-AHI communities without a significant relationship between distance and *Ae. aegypti* were Bretaña and Canelos. In Bretaña (AHI of 21.2%), there were two focal areas of presence, near the port and town center. We believe that the cluster near town center may be the only example of egg or immature-stage introduction of *Ae. aegypti* among our sites. Canelos (AHI 23.3%) had a marginal relationship between distance and *Ae. aegypti* presence. The town is slightly set back from the river and the *Ae. aegypti-*positive houses were in the half of town closer to the port, beginning just after the houses nearest to the port and ending about halfway through town. It is unclear whether this invasion occurred via adult or egg/immature life stages.

Given the global invasion of *Ae. aegypti,* there is surprisingly limited information regarding the life stage of invasion. Invasions have been associated with transit routes, but it is often unclear whether the invasion occurred via imported eggs, larvae, or adults [22, 42, 43]. In the case of the secondary dengue vector, *Ae. albopictus,* there is abundant evidence of the role of eggs and larvae as the invading life stages, and to a lesser extent, adults via passive transport in ground vehicles [44–46]. The evidence for passive dispersal of adult mosquitoes via boat presented by this study is unique in its level of detail and replication across sites.

The community-level distribution patterns also show that *Ae. aegypti* is at varying stages of invasion across different communities. In some locations it is well-established, while in others, it appears to be just beginning its invasion near port. Alternatively, this distribution pattern may suggest a failed invasion, in which the mosquito has not been able to advance further into the community. However, this seems unlikely because, anecdotally, the towns do not have internal expansion barriers. The identification of these active invasions provides a rare opportunity to monitor and understand the invasion process in real-time.

We found that the number of *Ae. aegypti* was significantly higher in houses with a mosquito larval habitat compared to those without. Similarly, the probability of adult *Ae. aegypti* presence was higher in houses with a larval habitat. These relationships existed despite not identifying mosquito larvae to species. Few other species in the region use containers as larval habitat and, based off a proportion raised to adulthood and *in-situ* morphological identification of larvae by eye, most larvae were likely *Ae. aegypti*.

Furthermore, the presence of a larval habitat, regardless of species, suggests that the container management of the household is permissive of mosquito breeding. The strength of these two relationships indicates that, in this context, the personal management of a household’s containers has a significant influence on that household’s exposure to adult *Ae. aegypti.* This highlights the importance of public health communication to encourage larval habitat removal. However, many houses without larval habitats still had adult *Ae. aegypti,* indicating that individual actions are not sufficient and must be supplemented by community-wide efforts. This aligns with findings of previous studies that reported weak or non-existent relationships between larval indices and adult indices, suggesting that larval indices are not the best predictor of adult *Ae. aegypti* exposure when measured in more detail [47–49].

The widespread presence of *Ae. aegypti* in rural Amazonia has significant public health implications. Over one million people live in rural communities in the Peruvian Amazon [36, 37, 50–52]. With the arrival of *Ae. aegypti,* these rural communities are newly at risk for dengue, Zika, and chikungunya viruses. Many of the communities are located far from hospitals [25, 26], with some study sites over 18 hours away by fast boat, which travels intermittently. Communities along less transited rivers face even greater access challenges. As a result of this inaccessibility, severe dengue cases that require immediate hospital care are evacuated by plane, a time-consuming and expensive effort. Early detection, timely hospitalization, and adequate care are critical for survival of hemorrhagic patients, all of which are limited in rural areas [53]. The expanding threat of dengue in the rural communities could result in a disproportionate burden of severe disease compared to urban areas due to the inaccessibility of care.

The widespread invasion of *Ae. aegypti* to rural Amazonia also increases the risk of sylvatic yellow fever virus (YFV) entering an urban transmission cycle. A large YF outbreak occurred across multiple regions in Peru in 1995 [54, 55], and between 2000 and 2014, Peru accounted for 37.4% of reported YF cases in the Americas [56]. Deforestation is hypothesized to increase YF outbreaks by bringing canopy-dwelling *Haemogogus* vectors to ground-level through tree felling, increasing mosquito-human contact [55]. This hypothesis was investigated in an ecological study, which failed to uncover a relationship between human activity (e.g. canopy tree loss) and YF case incidence, but the data may have been too coarse to accurately capture the relationship [56]. Recent Peruvian law changes permit more deforestation in the Amazon [57], which may increase YF transmission to people in remote villages, if deforestation is indeed a driver of human exposure. The presence of *Ae. aegypti* in rural areas increases the YFV amplification risk in rural human populations, thereby also increasing the probability of urban transmission by increasing the number of potential conduits to the city. This shifts the current understanding of the YFV rural transmission cycle as dependent on generalist mosquitoes [58], and may greatly expand amplification risk. A similar risk increase is also likely for other emerging arboviruses, such as Mayaro virus [59].

Our study shows an alarming level of *Ae. aegypti* expansion to rural Amazonia and highlights the need for attention to this new rural health threat. Given the public health risk demonstrated by our findings, we immediately alerted the regional health authorities, which has led to health department efforts to expand *Ae. aegypti* surveillance outside of the cities. However, this expanded surveillance will still be concentrated in larger towns due to limited resources.

There is still much to learn about this invasion process. For our study, we selected sites along the main shipping route; most are relatively large compared to other villages in the region, with a higher degree of fluvial transit connection to the cities. These characteristics may make our sites more likely to be invaded by *Ae. aegypti* than smaller, less connected communities. Future studies should focus on smaller, less connected communities to determine if certain community-level characteristics influence *Ae. aegypti* establishment. There should also be investigation into the transmission of dengue in rural communities. And finally, there is an urgent need to identify strategies to control rural *Ae. aegypti* populations in the region and stop further expansion of the species. Our research suggests that there is still time to intervene before the mosquito becomes fully established in many rural communities in the region.

## Acknowledgements

Thank you to everyone who made this research possible, especially the communities who welcomed us into their homes and allowed us to collect in their houses. Thank you to Drs. Gissella Vasquez and Ryan Larson (NAMRU SOUTH) for their support for this project since the early days of its conception, including permits and study design, and whose authorship on this publication is awaiting institutional clearance by NAMRU SOUTH. Thank you to Dr. Erika Mudrak (Cornell Statistical Consulting Unit) for advice on the statistical analysis, Pilar Díaz (GERESA Loreto) for providing data on the historical insecticide-use by GERESA in the communities, and to Gabriela Vásquez La Torre (Prisma) and Dr. Helvio Astete (NAMRU-SOUTH) for logistical support and advice. Our sincere gratitude to the Cornell Atkinson Center for Sustainability for funding support through the Rapid Response and Academic Venture Funds.

